# Characterization of the interaction between SARS-CoV-2 Membrane Protein and Proliferating Cell Nuclear Antigen (PCNA) as a Potential Therapeutic Target

**DOI:** 10.1101/2021.12.06.471464

**Authors:** Érika Pereira Zambalde, Isadora Carolina Betim Pavan, Mariana Camargo Silva Mancini, Matheus Brandemarte Severino, Orlando Bonito Scudero, Ana Paula Morelli, Mariene Ribeiro Amorim, Karina Bispo dos Santos, Mariana Marcela Góis, Daniel Augusto de Toledo-Teixeira, Pierina Lorencini Parise, Thais Mauad, Marisa Dolhnikoff, Paulo Hilário Nascimento Saldiva, Henrique Marques-Souza, José Luiz Proenca-Modena, Armando Morais Ventura, Fernando Moreira Simabuco

## Abstract

SARS-CoV-2 is an emerging virus from the *Coronaviridae* family and is responsible for the ongoing COVID-19 pandemic. In this work, we explored the previously reported SARS-CoV-2 structural membrane protein (M) interaction with human Proliferating Cell Nuclear Antigen (PCNA). The M protein is responsible for maintaining virion shape, and PCNA is a marker of DNA damage which is essential for DNA replication and repair. We validated the M PCNA interaction through immunoprecipitation, immunofluorescence co-localization, and a PLA assay. In cells infected with SARS-CoV-2 or transfected with M protein, using immunofluorescence and cell fractioning, we documented a reallocation of PCNA from the nucleus to the cytoplasm and the increase of PCNA and γH2AX (another DNA damage marker) expression. We also observed an increase of PCNA and γH2AX expression in the lung of a COVID-19 patient by immunohistochemistry. In addition, the inhibition of PCNA translocation by PCNA I1 and Verdinexor led to a reduction of plaque formation in an in vitro assay. We, therefore, propose that the transport of PCNA to the cytoplasm and its association with M could be a virus strategy to manipulate cell functions and may be considered a target for COVID-19 therapy.

## Introduction

The COVID-19 was claimed a global public health emergency by the World Health Organization (WHO). By November 17^th^, 2021, a total of 254.256.432 cases were confirmed, with more than five million deaths worldwide ^1^. COVID-19 is caused by SARS-CoV-2, an emerging virus from the *Coronaviridae* family of positive single-stranded RNA genome that encodes over 28 proteins, including 16 non-structural proteins (NSP1-NSP16), four structural proteins (spike, membrane, envelope, and nucleocapsid), and eight auxiliary proteins (ORF3a, ORF3b, ORF6, ORF7a, ORF7b, ORF8, ORF9b, and ORF14) ^2–4^. The four main structural proteins of the virion are responsible for: cell receptor recognition - spike (S); viral RNA packaging - nucleocapsid (N); and virus assembly - envelope (E) and membrane (M) proteins ^5–7^.

Nearly two years after the start of the pandemic, no treatment has yet been successful, and the infection mechanisms are still an open question ^8^. Many studies have been dedicated to understanding the interactome of SARS-CoV-2 with infected cells ^2,9–11^. The interaction between the structural protein M of SARS-CoV-2 and the human protein Proliferating Cell Nuclear Antigen (PCNA) was described with a Significance Analysis of INTeractome (SAINT) score of 1.0 ^2^, indicating a high probability of interaction. In addition, Stukalov et al. (2021) showed increased ubiquitination in specific regions (K13, K14, K77, K80, K248, and K254) of PCNA in cells infected with SARS-CoV-2 when compared to a control group.

M is the most abundant structural protein in the SARS-CoV-2 particle and is highly expressed ^12–14^. M is responsible for giving and maintaining the virion’s shape ^15^, is a three-pass membrane protein with three transmembrane domains and co-localizes with the endoplasmic reticulum (ER), Golgi apparatus, and mitochondrial markers ^15–17^. In addition, the M protein can interact with different coronavirus (CoVs) proteins: the N protein, which helps in viral genome packing ^18–20^, and the S protein, for its retention in the ER-Golgi intermediate compartment and its integration into new virions ^21^.

It was demonstrated that, in some CoVs, the M protein is highly immunogenic and induces specific T-cell responses after infection ^22^. In SARS-CoV-2 infection, M is also targeted by the immune response and plays a critical role in virus-specific B-cell response due to its ability to induce the production of efficient neutralizing antibodies in Covid-19 patients ^13,23^. During SARS-CoV-2 infection, the M protein can directly bind to MAVS (Mitochondrial Antiviral Signaling Protein) to inhibit the innate immune response ^24^. More specifically, the SARS-CoV-2 M protein antagonizes type I and III IFN production by targeting RIG-I/MDA-5 signaling and preventing the multiprotein complex formation of RIG-I, MAVS, TRAF3, and TBK1. This multiprotein complex blocks the activation of IRF3-induced anti-viral immune suppression, facilitating virus replication ^17,24^. Although the M protein binding to MAVS mechanism has been described, other authors suggest that M is mainly involved in ATP biosynthesis and metabolic processes ^9,25^, indicating that it has different functions.

The PCNA is a 36 kDa protein, well-conserved in all eukaryotic species, from yeast to humans ^26^. This protein controls essential cellular processes such as DNA replication and damage repair, transcription, chromosome segregation, and cell-cycle progression ^26–31^. It has been dubbed the “maestro of the replication fork” ^32^. PCNA, as a homotrimer, encircles duplex DNA, forming a ring-shaped clamp ^33^. The PCNA is a scaffolding protein that interacts with several other proteins, mainly involved in DNA replication and repair ^31^. In most cell types, PCNA is exclusively nuclear, but studies demonstrated that it could go to the cytoplasm. In cancer cells, cytoplasmic PCNA was described as a regulator of the cell metabolism binding to enzymes involved in the glycolysis pathway, regulation of the energy-generating system in mitochondria, cytoskeleton integrity, and other cellular signaling pathways through binding to cytoplasmic and membrane proteins ^34^. PCNA is cytosolic in mature neutrophils and acts in immune response, including to virus infection ^35–38^.

Although many studies indicate the PCNA role in DNA virus infection, its association with RNA viruses is poorly understood. To our knowledge, the only study that observed a function of PCNA in an RNA virus was with the Bamboo Mosaic virus (BaMV), a common virus in plants. In this study, the authors demonstrated that PCNA goes to the cytoplasm and directly binds to the BaMV replication complex, downregulating the replication efficiency of the virus ^39^.

In this study, we validated the M-PCNA interaction and demonstrated that the M protein facilitates the transport of PCNA from the nucleus to the cytoplasm. M expression and SARS-CoV-2 infection were associated with increased phosphorylation of H2AX and increase of PCNA expression. Drugs that block PCNA translocation from nucleus to cytoplasm inhibit virus replication. This indicates a new mechanism in SARS-CoV-2 replication and a potential target for therapy.

## Results

### SARS-CoV-2 structural proteins interact with PCNA in HEK293T cells

In the study by Gordon et al. (2020), the interactome of the structural and non-structural proteins of SARS-CoV-2 was performed in HEK293T cells ^2^. The analysis of affinity-purification–mass spectrometry (AP-MS)-based proteomics showed that PCNA interacts with E and M SARS-CoV-2 proteins. In our study, we validated this interaction through an anti-FLAG immunoprecipitation assay (Figure 1A). FLAG-tagged GFP, E, M, and N proteins were expressed in HEK293T cells and immunoprecipitated with anti-FLAG antibodies. As a result, we identified that PCNA co-immunoprecipitated with the E and M SARS-CoV-2 proteins but not with the N protein, which is expected since PCNA was not identified as an interactor of N protein ^2^. Anti-FLAG immunoprecipitation confirmed the interaction between M and PCNA in two independent experiments (Supplementary Figure S1). We also confirmed that PCNA and M interact, through a reverse immunoprecipitation assay (Figure 1B), in HEK293T cells previously transfected with FLAG-tagged GFP, E, M, and N. We identified that only FLAG-tagged M co-immunoprecipitated with PCNA, revealing a more specific interaction between these proteins. Furthermore, we explored the interaction between FLAG-M and PCNA by proximity ligation assay (PLA). Vero E6 cells expressing FLAG-M were fixed 24 h.p.t. (hours post-transfection) and labeled with primary antibodies against FLAG (rabbit) and PCNA (mouse), followed by PLA probes conjugated to anti-rabbit or anti-mouse. PLA signal is emitted when probes attached to primary antibodies are closer than 40 nm, indicating protein interaction. As negative controls, transfected cells were labeled with only one of the primary antibodies or omitting both. Figure 1C (panels A-C and supplementary figure S2) shows positive PLA dots in transfected cells, while minimal or no signal is seen in the respective controls (Figure 1C – panels D-L). A confocal microscopy analysis of the PLA assay also showed that the M-PCNA interaction occurs in the cytoplasm, indicating a possible translocation of the PCNA to the cytoplasm, induced by the M protein (supplementary Figure S3).

**Figure 1.**
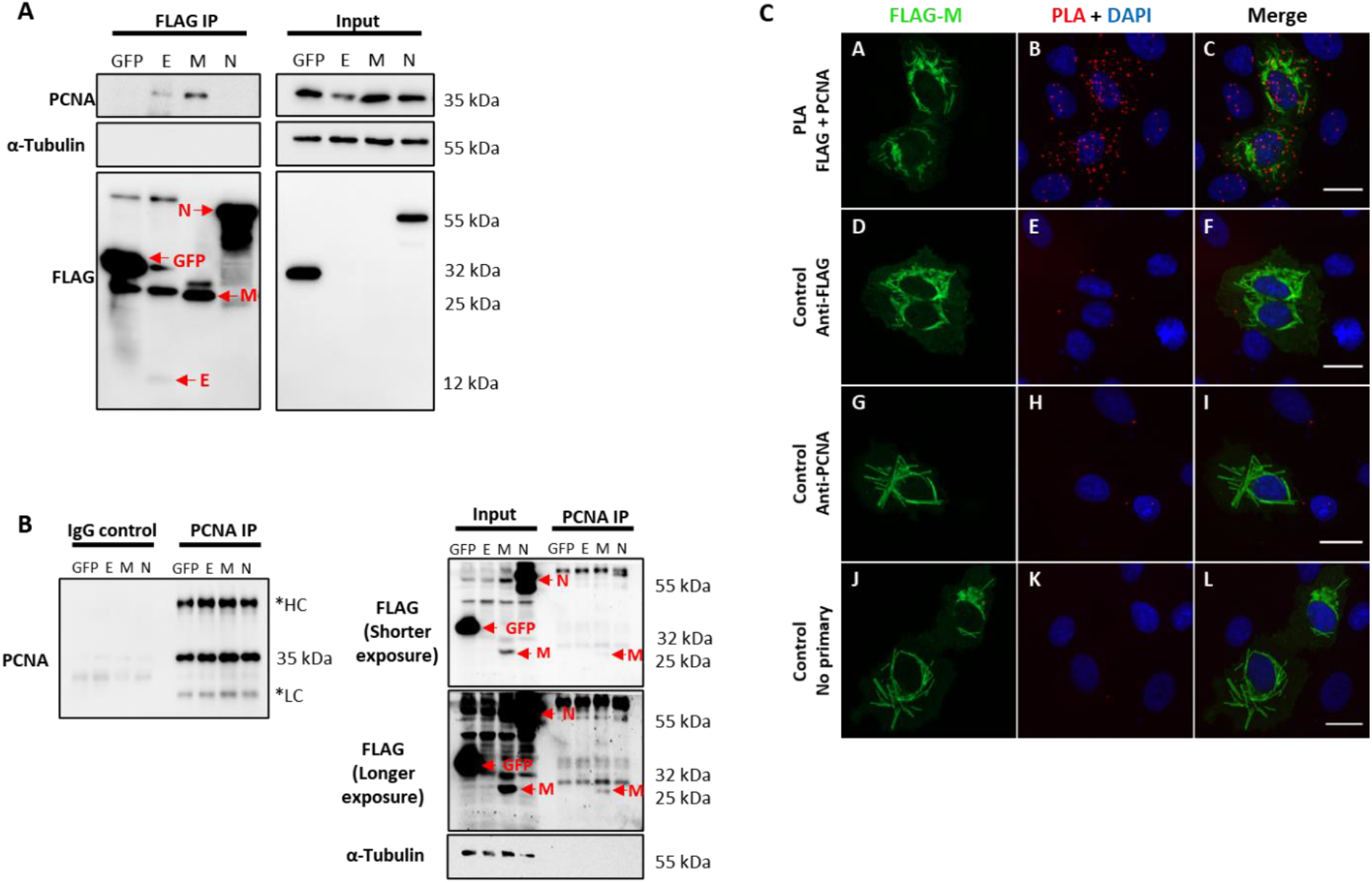
SARS-CoV-2 structural proteins interact with PCNA in HEK293T and Vero E6 cells. FLAG-tagged GFP, E, M, and N proteins (indicated at the top of the panel) were expressed by transfection in HEK293T cells. Immunoprecipitation with anti-FLAG (A) or anti-PCNA antibodies (B) was performed (indicated on panel’s top). Blots were done with the primary antibodies indicated on the panel’s left, and Molecular Weight markers sizes are indicated on the right. A) PCNA was identified as an interactor of SARS-CoV-2 E and M proteins. B) Immunoprecipitation of endogenous PCNA confirmed that protein M co-immunoprecipitated with PCNA. These data were obtained in one biological replicate. * HC: Heavy Chain, LC: Light chain. Red arrows indicate specific bands. C) Vero E6 cells were transfected with pFLAG-M and submitted to ice-cold methanol fixation after 24h, followed by incubation with the primary antibodies, as indicated on the left, and PLA staining protocol. Panels A, B, and C show positive PLA dots for FLAG and PCNA staining in transfected cells. Little to no signal is detected in the respective controls omitting one or both primary antibodies (panels D–L). Images are representative of two independent experiments. Nuclei were stained with DAPI. All images were taken at 63× magnification with a ZEISS Axio Vert.A1 microscope.

### SARS-CoV-2 M expression in Vero E6 cells induces PCNA translocation to the cytoplasm

To better understand how the interaction of M-PCNA could be acting on the cells, we conducted immunofluorescence (IF) assays in Vero E6 cells expressing FLAG-M. 24 hours after transfection, cells were stained and analyzed by widefield microscopy (Figure 2A). Although PCNA did not entirely co-localizes to the structures where M is present, we observed that in transfected cells, PCNA presented a more cytosolic pattern when compared to the non-transfected ones, as indicated by plot profile analysis (Figure 2A – Arrows 1 and 2). To further investigate this phenomenon, we quantified the fluorescence intensities of nuclear and cytoplasmic PCNA in cells expressing FLAG-M and non-transfected cells. Figure 2B shows that in non-transfected cells, PCNA has a propensity to be nuclear, but under FLAG-M expression, the PCNA translocated to the cytoplasm. We also evaluated the translocation of PCNA to the cytoplasm by odds ratio. The odds ratio for cytoplasmic PCNA with FLAG-M expression was 6.34 (Confidence Interval 95% =2.83-13.3) compared to non-transfected cells, indicating the correlation of M protein with the translocation of the PCNA to the cytoplasm. In addition, the subcellular fractionation of Vero E6 cells transfected with FLAG-M or FLAG-GFP was carried out (Figure 2C). In agreement with IF data, FLAG-M transfected cells showed an evident reduction of PCNA in the nuclear fraction, while it remained nuclear in cells expressing FLAG-GFP (Figure 2C). These results indicate that M protein acts on PCNA translocation from the nucleus to the cytoplasm.

**Figure 2.**
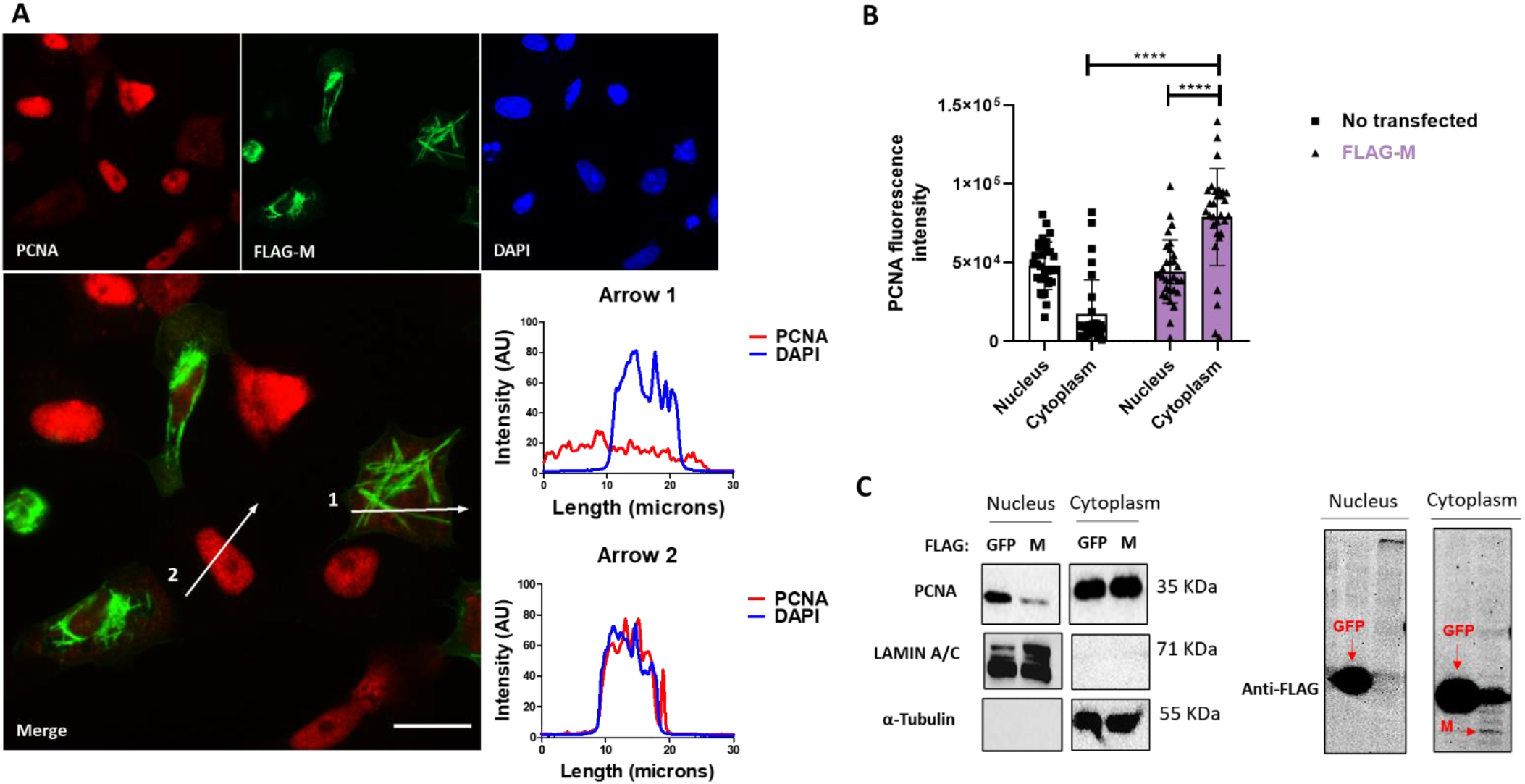
FLAG-M expression promotes PCNA translocation to cytoplasm. A) Vero E6 cells expressing FLAG-M protein, 24h after transfection, were fixed and stained for FLAG-M (green), PCNA (red), and DAPI. Graphs show plot profile intensities for PCNA and DAPI channels in the cross-sections indicated by arrows 1 and 2. Images are representative of two independent experiments. Scale bar 20 μm. B) PCNA localization was analyzed through fluorescence intensity in non-transfected and FLAG-M transfected cells. Briefly, nuclei area stained with DAPI was selected manually and the ROI (region of interest) obtained was used to measure PCNA intensities on nucleus and cytoplasm. Fluorescence intensity was quantified in grayscale on ImageJ. This data is representative of two independent experiments. The data represent mean ± SD (n=30). For statistical analysis, Two-way ANOVA and multiple comparison Bonferroni’s tests were used. *p<0.05 and **** p<0.0001 were considered statistically significant. C) Vero E6 cells were transfected with vectors to express FLAG-tagged M or GFP proteins, and after 48 hours cellular fractionation followed by Western blotting was performed. LAMIN A1 and α-Tubulin proteins were used as controls for nuclear and cytosolic fractions, respectively. The expression of the transfected proteins is shown in the right panel. Western blot images are representative of one independent experiment.

### SARS-CoV-2 M protein interacts with PCNA in the cytoplasm of Vero E6 cells

To confirm the interaction between M and PCNA, we performed confocal immunofluorescence of FLAG-M transfected cells. Plot profile analysis shows an overlap between FLAG-M and PCNA signals, indicating the proximity of the proteins (Figure 3). Moreover, as mentioned before, interaction between FLAG-M and PCNA was detected by PLA (Figure 1C), and the co-localization of PLA puncta with FLAG-M was shown by confocal microscopy (Supplementary Figure S3). Taken together, these results indicate the ability of M protein in inducing PCNA translocation from the nucleus, and to interact with it in the cytoplasm.

**Figure 3.**
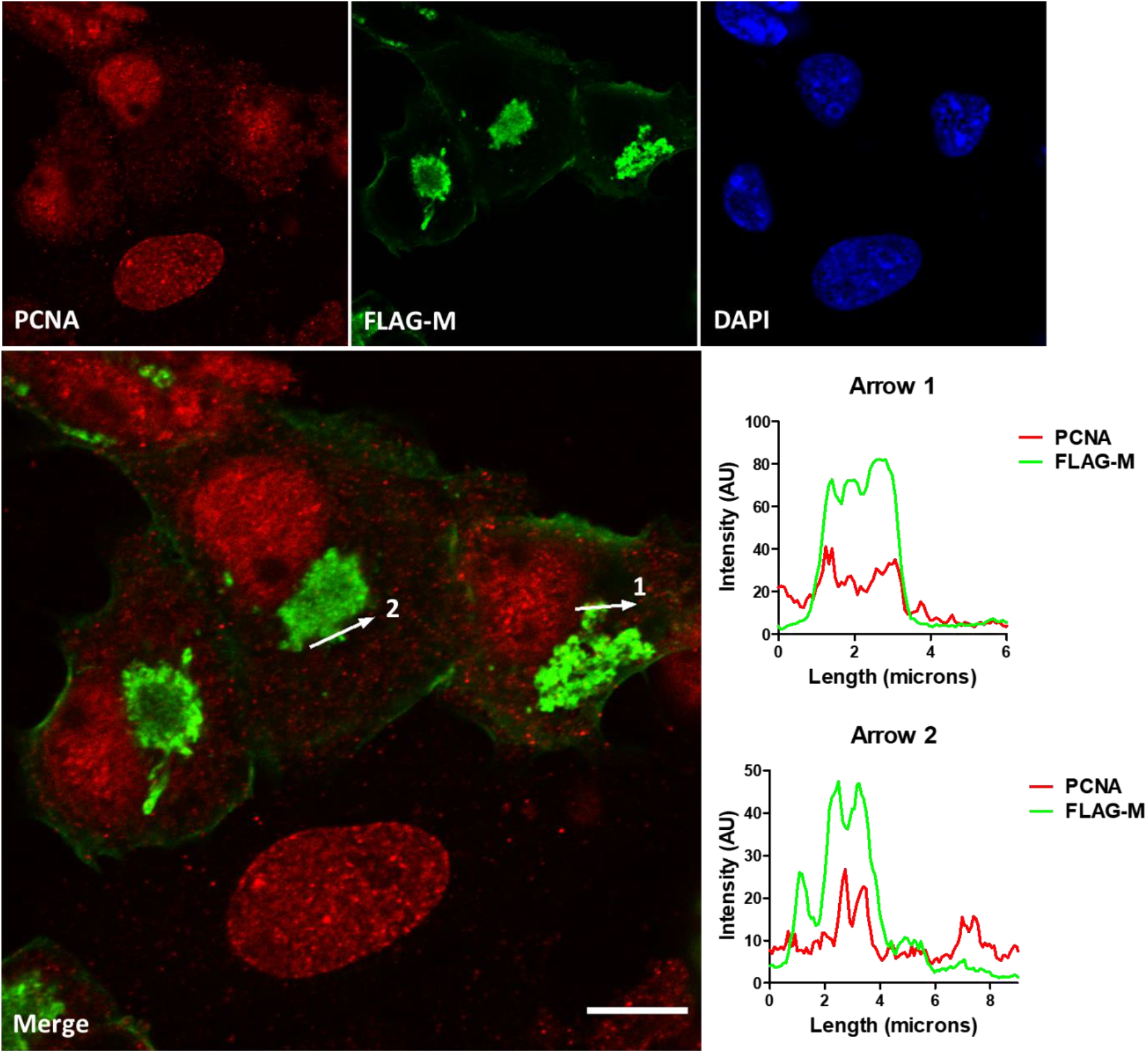
Co-localization of FLAG-M and PCNA by confocal immunofluorescence. Vero E6 cells expressing FLAG-M and stained for PCNA (red) and FLAG-M (green) were analyzed by confocal microscopy. The plot profile of two arrowed areas (arrows 1 and 2) indicates overlap in signals of both proteins, as shown in the graphs on the right. Images were taken at 100× magnification with a Zeiss LSM-780-NLO microscope. Scale bar 10 μm.

### FLAG-M expression increases DNA damage marker levels in HEK293T and Vero E6 cells

PCNA is one of the essential proteins for DNA replication and DNA repair, and it is also known as a DNA damage marker. To address whether M expression could be related to the induction of DNA damage. We transfected HEK293T and Vero E6 cells and looked at γH2AX and PCNA expression levels since both proteins elevated expression is associated with DNA damage. In HEK293T transfected cells, we could observe a slight increase in the levels of PCNA and γH2AX (Figure 4A), but no significant difference was observed in the Vero E6 cell line (Figure 4B). Because transfection efficiency could be a limiting factor for detecting inconspicuous events like DNA damage, we also explored γH2AX levels in transfected Vero E6 cells by immunofluorescence (Figure 4C). Comparing the γH2AX fluorescence intensity in FLAG-M transfected cells with non-transfected cells, we found a significant increase in the phosphorylation level of H2AX, suggesting that M involvement with PCNA may promote DNA damage (Figure 4D).

**Figure 4.**
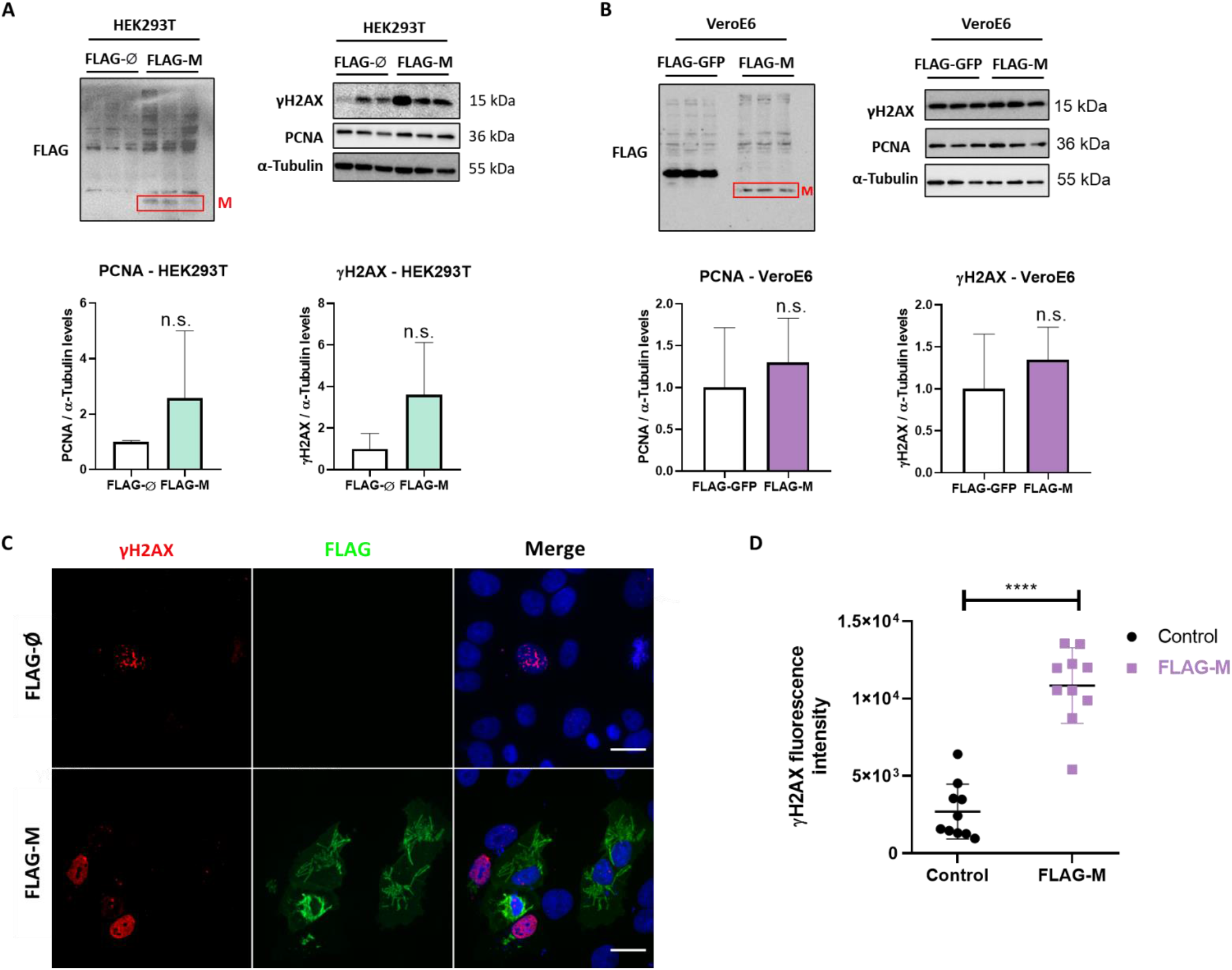
PCNA and γH2AX levels in transfected cell lines. A) Detection of PCNA and γH2AX levels in HEK293T cells transfected with pFLAG-M compared to pFLAG (empty vector). B) Detection of PCNA and γH2AX levels in Vero E6 cells transfected with pFLAG-M compared to pFLAG-GFP. Graphs show a slight increase in normalized protein levels; however, the statistical test does not show significance. Western blotting data in A) and B) are representative of one independent experiment. C) Immunofluorescent staining of FLAG-M (green) and γH2AX (red) in Vero E6 cells 24 hours post-transfection. Images are representative of two independent experiments. Scale bar 20 μm. D) Fluorescence intensity quantification of γH2AX levels in FLAG-M transfected versus non-transfected cells. Fluorescence intensities on nucleus were measured as described in Figure 2B. Data represent mean ± SD in samples from 2 independent experiments (n = 10). For statistical analysis, a two-tailed unpaired T-test was conducted. n.s. non-significant, **** p<0.0001 was considered statistically significant.

### SARS-CoV-2 infection induces PCNA translocation to the cytoplasm and promotes DNA damage

To verify if our findings concerning M protein and the DNA damage markers PCNA and γH2AX were also reproducible during infection, we evaluated the effect of SARS-CoV-2 infection in Vero E6 cells. Immunofluorescence analysis showed PCNA staining in the cytoplasm of infected cells, with PCNA presenting mainly a nuclear pattern on mock control (Figure 5A). The results (Figures 5B and C) show a higher intensity of PCNA in the cytoplasm of infected cells with a concomitant reduction in its nuclear signal compared to mock-infected cells. In addition, a higher γH2AX fluorescence intensity was observed in infected cells compared to non-infected cells, which confirms that the infection promotes DNA damage (Figure 5D and E). We also evaluated PCNA and γH2AX levels by western blotting in HEK293T and Vero E6 infected cells (Figure 5F). The effect observed during infection is a higher expression of PCNA and phosphorylation of histone H2AX in both cell lines (Figure 5G and H).

**Figure 5.**
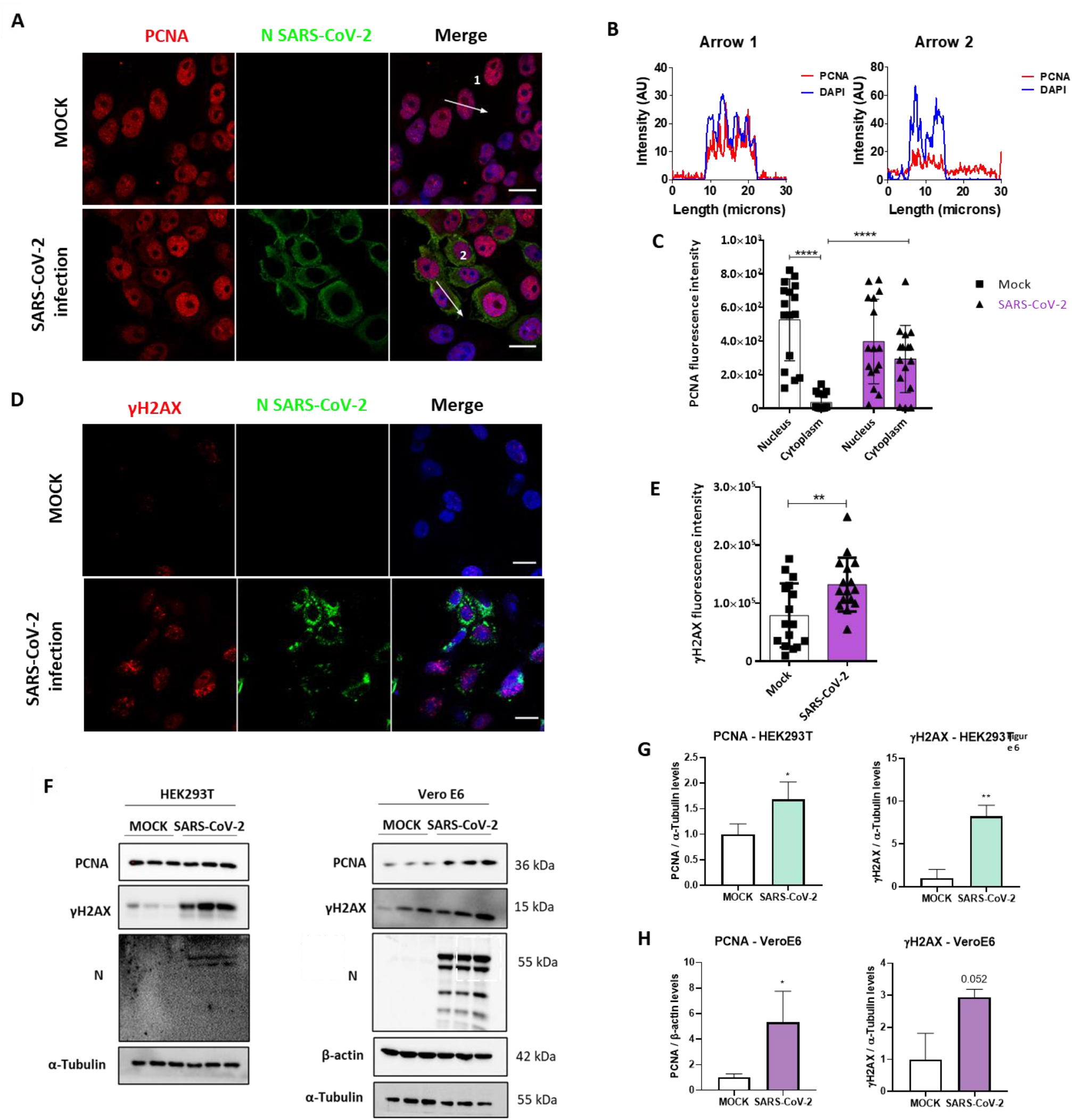
SARS-CoV-2 infection promotes PCNA translocation to the cytoplasm and enhances PCNA and γH2AX expression. A) Vero E6 cells were infected with SARS-CoV2 (MOI 0.3), and 24 hours post-infection immunofluorescence was performed for N (green) and PCNA (red). Scale bars 20 μm. B) Plot profile intensities for PCNA and DAPI channels in the cross-sections indicated by arrows 1 and 2. Images are representative of two independent experiments. Scale bar 20 μm. C) PCNA localization was analyzed through fluorescence intensity mock versus infected cells, as described in Figure 2B. White bars = mock, purple bars = SARS-CoV-2 infection. D) Vero E6 cells were infected with SARS-CoV2 (MOI 0.3), and 24 hours post-infection immunofluorescence was performed for N (green) and γH2AX (red). Scale bars 20 μm. E) Fluorescence intensity of γH2AX in the nucleus was analyzed in mock and infected cells, as described in Figure 2B. White bars = mock, purple bars = SARS-CoV-2 infection. F) Western blotting analysis of PCNA and γH2AX levels in HEK293T and Vero E6 cells infected with SARS-CoV-2 compared to mock. Statistical analysis for normalized expression levels of PCNA and γH2AX are shown for HEK293T (G) and Vero E6 cells (H). Data represent means ± SD from 1 independent experiment. For fluorescence intensity (n = 20), data was analyzed by Two-way ANOVA and multiple comparisons Bonferroni’s test. *p<0.05 and ****p<0.0001 were considered statistically significant.

We then compared PCNA (top panels) and γH2AX (bottom panels) expression levels in lung sections obtained from control and COVID-19 patients through immunohistochemistry assay (Supplementary Figure S4). Our results indicate high expression of both markers in COVID-19 patient (right panels), suggesting the involvement of both PCNA and γH2AX in SARS-CoV-2 infection in vivo.

### Stabilization of PCNA trimer by PCNA-I1 or blockage of CRM-1 transporter by Verdinexor inhibits SARS-CoV-2 replication

Our data indicate that PCNA translocate to the cytoplasm after M protein expression or SARS-CoV2 infection. To analyze if this phenomenon plays a role in the replicative cycle of the virus, two inhibitors of this transport were used to prevent the PCNA migration to the cytoplasm and test their possible anti-viral activity. The inhibitor of PCNA (PCNA I1) prevents the transportation of the PCNA to the cytoplasm by stabilizing its trimer form in the nucleus. Verdinexor is a selective inhibitor of nuclear export protein exportin-1 (XPO1), also called Chromosome Region Maintenance 1(CRM1), and was already described as essential in the PCNA translocation ^36,40^. Vero E6 cells were infected and submitted to an anti-viral in vitro efficacy assay. The results (Figure 6) indicate that PCNA I1 0.5 μM or Verdinexor 0.1 μM present effects against SARS-CoV-2, reducing the viral replication roughly by 20% and 15%, respectively, compared to the DMSO.

**Figure 6.**
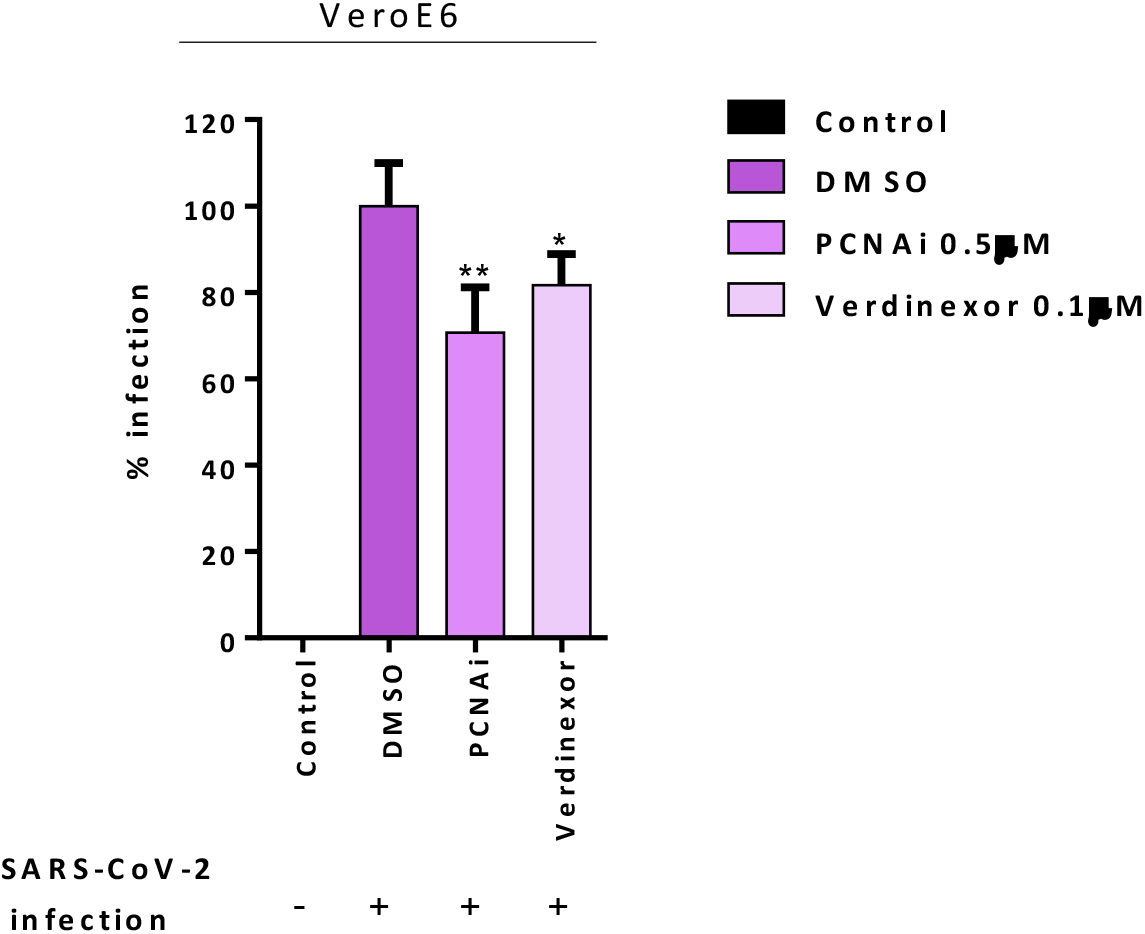
PCNA I1 and Verdinexor inhibit SARS-CoV-2 viral replication in vitro. Vero E6 cells were infected with SARS-CoV-2 as described in anti-viral in vitro efficacy assay (see Methods), for one hour, then DMSO, PCNA I1 0.5 μM or Verdinexor 0.1 μM were added to overlay media one hour after virus adsorption. The viral load was assessed by plaque assay after four days of incubation. This data is representative of two independent experiments. T-test was used for independent comparisons between DMSO versus PCNA I1, and DMSO versus Verdinexor. *p<0.05 and **p<0.001 were considered statistically significant.

## Discussion

After nearly two years since the start of the COVID-19 pandemic, understanding the underlying mechanisms of SARS-CoV-2 infection and the search for treatment are still in high demand. Large-scale analysis of SARS-CoV-2 human infection interactome indicated several protein-protein interactions that require further validation ^9–11,43^. In this work we aimed to characterize the interactions of viral proteins and PCNA. According to Gordon et al. (2020) ^2^, the score of SAINT probability of PCNA for E protein was zero (a low score of interaction probability), and for M was 1.0 (a high score of interaction probability), while PCNA did not appear as an interactor for N protein. Through immunoprecipitation and PLA assays (Figure 1), we validated the M-PCNA interaction. The E-PCNA interaction was not detected in the reverse immunoprecipitation (PCNA pull down, Figure 1B), and we focused on the M-PCNA interaction.

PLA assay indicates that M-PCNA interaction occurs in the cytoplasm (Figure 1C), and to better characterize it, we performed a co-localization assay by immunofluorescence and subcellular fractioning. Our data indicate that in M expression (Figure 2) or upon SARS-CoV-2 infection (Figures 5 A, B, C), PCNA translocate from nucleus to cytoplasm. In agreement with PLA data, upon M expression there is also a partial co-localization among M and PCNA in the cytoplasm documented by confocal microscopy (Figure 3 and Supplementary Figure S3). This can be hypothesized as a need for the virus to have PCNA partially translocated from the nucleus to the cytoplasm, and that M is responsible, at least in part, for this translocation in the context of the viral infection.

PCNA acts as a co-factor for DNA polymerases in normal conditions, being essential for DNA replication ^44,45^. PCNA also participates in DNA repair, on the metabolism of DNA, and chromatin by recruiting various enzymes, and not only by acting as a scaffold and localizing these factors, but also activating their enzymatic activities ^46^. Aside from this, post-translational modifications in PCNA alter its function in different ways ^32,46,47^. The interactome described by Stukalov et al. (2021) demonstrated that higher ubiquitination occurs in specific regions of PCNA in SARS-CoV-2 infection, indicating a regulatory mechanism of PCNA by the virus infection ^10^.

In cells transfected with Flag-M (Figure 4) or infected with SARS-CoV-2, we observed an increase of PCNA and γH2AX (Figures 5 D to H), and both proteins are associated with DNA repair. We also found a high expression of PCNA and γH2AX in a COVID-19 patient (Figure S4). Li et al. (2021) already reported a high PCNA expression in moderate COVID-19 patients compared to the control group by proteomics analysis ^9^. It has been reported that virus infection in human cells can cause DNA damage, inhibiting the association of the DNA polymerase to the DNA stalling and collapsing the replication forks, which results in DNA double-strand breaks (DSB) ^48,49^. This damage activates a stress response, mediated by checkpoint kinases, which help stabilize and restart the replication forks, thus preventing the generation of DNA damage and genomic instability ^50^. In this case, one of the strategies of the cells is to trigger the PCNA ubiquitination. When PCNA is polyubiquitinated, it searches for damaged DNA to assemble the replication complex, with a less specific DNA polymerase ^51^. This mechanism is known as translesion synthesis (TLS) and is activated to bypass damaged DNA ^49^. Thus, the increase of PCNA could be a strategy for the viral infection to maintain cell viability. Also, after DNA damage, several proteins are activated to manage the DNA lesion. One of them is γH2AX, the phosphorylated form of H2AX responsible for recruiting repair proteins to deal with stalled forks ^52^, and also increases the expression of p53 and phosphorylation of p53, leading to cell growth inhibition ^53,54^.

PCNA is also found in the cytoplasm of cancer cells. The accumulation of PCNA in the cytoplasm of cancer cells evidenced interactions between PCNA and the proteins in the cytoplasm ^55^. PCNA can bind to enzymes of the glycolysis pathway and regulate energy production in the mitochondria; maintain cytoskeleton integrity; and participate in other signaling pathways ^11,12^. Li et al. (2021) described that SARS-CoV-2 M protein has its activity connected to ATP biosynthesis and metabolic processes ^9^. Considering these data, the PCNA translocation from nucleus to cytoplasm reported here, may also be associated with the regulation of metabolism in infected cells to maintain SARS-CoV-2 replication.

Another hypothesis is that cytosolic PCNA, in association with the M protein, could bind to proteins that inhibit the immune response. Several studies pointed out that cytoplasmic PCNA is found in mature neutrophils associated with procaspases, thus preventing neutrophils from apoptosis ^37^. The accumulation of PCNA in the cytoplasm of mature neutrophils is due to the activity of the chromosome region maintenance 1 (CRM1)-dependent nuclear-to-cytoplasmic relocalization during granulocytic ^36^. Cytoplasmic PCNA is also associated with Caspase-9 in the SH-SY5Y neuroblastoma cell line, and S-nitrosylation of PCNA at the residues C81 and C162 decreases this interaction, leading to caspase-9 activation ^56^.

Verdinexor is a selective inhibitor of nuclear export (SINE), a molecular drug that binds to CRM-1 and blocks the transport of proteins from the nucleus to the cytoplasm, including PCNA ^36,57^. The CRM-1 inhibitors have demonstrated activity against over 20 different DNA and RNA viruses, including influenza and respiratory syncytial virus ^57–59^. In addition, Selinexor, another SINE, reduced SARS-CoV-2 infection in vitro and protected the respiratory system in an in vivo model ^42^. We found that Verdinexor reduced viral replication by 15% (Figure 6), corroborating the previous SINE study. Nonetheless, SINEs are drugs that could act in the translocation of different proteins, not only in PCNA. We also treated cells with a PCNA inhibitor (PCANI1). This inhibitor stabilizes the PCNA trimer and prevents the PCNA translocation from the nucleus to the cytoplasm and reallocation inside the nucleus^41,50^. This inhibits the PCNA action on the replication forks and indirectly leads tumor cells to a higher sensitivity to anticancer DNA damaging drugs ^6,14^. Our results show that PCNA I1 reduced viral replication by 20% (Figure 6), indicating a potential use of PCNA translocation inhibition as a treatment for COVID-19.

## Conclusions

The SARS-CoV-2 virus can manipulate many pathways to replicate in the host cell. In this study, we validated the PCNA-M interaction and underlined some of the potential mechanisms regarding this interaction. The increase of the PCNA and γH2AX levels in the nucleus may prolong cell viability to favor virus replication. On the other hand, the translocation of PCNA and its association with M in the cytoplasm may manipulate the immune response and regulate cell metabolism to favor virus replication. Finally, inhibition of PCNA translocation from the nucleus to cytoplasm reduced the formation of plaques in an in vitro assay, indicating a potential therapeutic target. The original data reported here may help to better understand the SARS-CoV-2 replication mechanisms, thus impacting therapeutic strategies for COVID-19 resolution.

## Methods

### Cell culture

The VeroE6 (African green monkey, *Cercopithicus aethiops*, kidney) and HEK293T (Human embryonic kidney) cell lines were cultivated in Dulbecco’s modified Eagle (DMEM) (Thermo Scientific #12100-046) medium, supplemented with 10 % fetal bovine serum (FBS # 12657029) and 1 % penicillin/streptomycin (Gibco #15140-122). Cells were maintained at 37 °C in a humidified atmosphere containing 5 % carbon dioxide.

### Viral Infection

An aliquot of SARS-CoV-2 SP02.2020 (GenBank accession number MT126808) isolate was kindly donated by Dr. Edison Luiz Durigon (Institute of Biomedical Sciences, University of São Paulo). Vero E6 cells were used for virus propagation in the Biosafety Level 3 Laboratory (BSL-3) of the Laboratory of Emerging Viruses (Institute of Biology, State University of Campinas). Viral infections were performed in Vero cells seeded in 24-well plates (5 × 10^5^ cells/well) for the experiments with treatments and immunofluorescence assays, and six-well plates (1 × 10^6^ cells/well) for Western blots. A multiplicity of infection (MOI) of 0.3 was used for all experiments.

### Cloning

To do the FLAG-tagged protein expression, a modified pcDNA 3.1 (+) (Thermo Fisher Scientific) was generated by cloning the FLAG peptide coding sequence upstream of the multiple cloning sites, using the *Nhe*I and *Bam*HI restriction sites and generating the pcDNA-FLAG vector ^60^. The full-length of M, N, and E genes were codon-optimized, synthesized (Geneart-Thermo Fisher Scientific), and cloned into pcDNA-FLAG, generating plasmid pFLAG-M, pFLAG-N, and pFLAG-E. The pFLAG-green fluorescent protein (GFP) ^61^ was used as the control plasmid for the expression of a non-related protein in the immunoprecipitation assays.

### Transfection

VeroE6 and HEK293T cells were seeded for 24 hours before transfection. Transfection was performed with Lipofectamine (Thermo Scientific - #20071882) and PLUS reagents (Thermo Scientific - #15338100). The protocol of plasmids’ transfection is described by Amaral et al 2016 ^61^. For immunofluorescence, the cells were seeded and transfected in 24-well plates; for western blotting in 6-well plates; and for immunoprecipitation in 100 mm plates.

### Immunoprecipitation

For anti-FLAG immunoprecipitation, HEK293T cells cultivated in 100 mm diameter dishes expressing the FLAG-tagged GFP, E, M, and N proteins were washed twice with PBS and harvested by pipetting up and down in PBS after 48 hours. Cells were resuspended in 500 μL of lysis buffer (50 mM Tris-HCl, pH 7.4, 150 mM NaCl, 1 mM EDTA, 1% Triton X-100) containing protease inhibitor cocktail (Roche). Protein lysates were incubated and shaken on ice for 15 min and centrifuged at 12,000 × g for 10 min at 4°C. Supernatants were collected and protein quantification was performed using a BCA protein assay kit (Thermo Scientific). A total of 2,000 μg of protein extract was used to perform the immunoprecipitation, so the samples were diluted with lysis buffer without inhibitors and incubated overnight at 4°C with 30 μL of anti-FLAG agarose-coupled beads (#A2220, Sigma-Aldrich) under mild agitation. Subsequently, the beads were washed five times with 600 μL of ice-cold TBS (50 mM Tris-HCl, pH 7.5, 150 mM NaCl) and eluted with 300 ng/L of FLAG peptide (#F4799, Sigma-Aldrich) for four hours under moderate agitation ^62^. Supernatants were collected and stored at −20°C for immunoblotting analysis.

For reverse immunoprecipitation, in HEK293T cells, the same protocol of protein lysis and quantification were performed. A total of 300 μg of protein extract was used to perform the immunoprecipitation. The samples were diluted with lysis buffer without inhibitors and incubated overnight at 4°C with anti-PCNA antibody (1:500, #2586, Cell signaling) under mild agitation. Subsequently, 30 μL of protein A/G Agarose (#20421, Thermo Scientific) were added to each sample, followed by mild agitation for two hours. Samples were washed five times with a wash buffer (25 mM Tris-HCl, pH 7.5, 150 mM NaCl) and the elution was performed with Laemmli Buffer with SDS 1×. Supernatants were collected and stored at −20°C for immunoblotting analysis.

### Immunofluorescence

Sterilized glass coverslips were treated with 6M HCL (Synth) and placed in each well on 24-well plates. Vero E6 cells were seeded at a density of 1×10^5^ cells in DMEM with 10% FBS. After the experimental procedures, the wells were washed 1 × with PBS, permeabilized with ice-cold methanol 100% for 10 minutes, or fixed with paraformaldehyde 4% (PFA – Sigma Aldrich 158127) for 15 minutes. The cells were then permeabilized with PBS-Tween-20 0.1% (PBS-T - Sigma Aldrich P1379) for 10 minutes, blocked with 1% BSA-Tween-20 0.3% for 30 minutes, and incubated overnight with a solution containing the primary antibody in (1:200) 1% BSA-Tween-20 0.3% at 4°C. After a washing step with PBS (three times for 10 minutes each), a solution containing the secondary antibody (1:250) and PBS-Tween20 0.1% was added to each well for one hour in a dark chamber. The wells were then washed three times with PBS (for 10 minutes each), a solution containing Hoechst (Sigma Aldrich – 861405 - 1:1,000) was added for 10 minutes, followed by three times washing with PBS (for 10 minutes each), the glass coverslips were removed from the wells with the aid of tweezers and added to glass slides with 5 μL of Glycerol (Sigma Aldrich G5516). The primary antibodies used were: PCNA (Cell signaling #2586), N (Invitrogen #MA5-35943), Flag (Sigma # F3165). The secondary antibodies used were: Alexa-Fluor-594 Goat anti-Rabbit (Jackson ImmunoResearch #711-585-152), Alexa-Fluor-488 Goat Anti-Mouse (Jackson ImmunoResearch #705-545-003). To quantify the signal on immunofluorescences we used the ImageJ software to transform the images in 8 bit (grayscale), and check-in Set Measurements the options: Area and Integrated density; select an ROI of nuclei area (Hoechst staining) to determinate the fluorescence signal on nuclei; the cytoplasmatic fluorescence signal was obtained by a total cell - nuclei area ROI. Besides that, we computed the signal of the background (region without cells). Finally, the Corrected Cell Total Fluorescence (CTCF) was calculated by multiplication of the integrated density of gray, of nuclei or cytoplasm by their respective areas and subtracting the background signal.

### Subcellular Fractionation

Vero E6 cells were seeded at 8×10^6^ in a P100 plate and transfected with pFLAG-GFP or pFLAG-M. After 48 hours, the subcellular fractionation was finalized (Baldwin, 1996) ^63^. Briefly, first the cytoplasmic fraction was isolated from the nuclear solution using a cytoplasmic buffer (10 mM HEPES; 60 mM KCl; 1 mM EDTA; 0.075% NP-40; 1 mM DTT; protease inhibitor 1×) (pH=7.6) and centrifuged in 1,400×g for 30 minutes. The cytoplasmic supernatants were collected into a new tube. The nuclear pellet was washed with cytoplasmic buffer without NP-40 two times followed by a five-minute centrifugation at 1,000×g. The final pellet was resuspended with nuclear buffer (20 mM TrisCl; 420 mM NaCl; 1.5 mM MgCl_2_; 0.2 mM EDTA; 25% glycerol; protease inhibitor 1×) (pH=8.0) and incubated on ice for 10 minutes, and vortexed periodically. Finally, the cytoplasmic and nuclear solutions were centrifuged at 12,000×g for 10 minutes and the supernatants were collected into new tubes. The quantification was performed by BCA.

### Proximity Ligation Assay (PLA.)

For PLA protocol, Vero E6 cells were grown on chambered slides at a confluency of 70% and transfected with pFLAG-M. After 24 hours, cells were fixed with ice-cold methanol for 15 min and permeabilized with 0.1% TritonX-100 in PBS. The next steps faithfully followed the manufacturer’s specifications (Duolink^®^ In Situ – Sigma-Aldrich). Briefly, cells were blocked and stained with primary antibodies (anti-FLAG 1:400 - Sigma F7425 and anti-PCNA 1:400 – Sta Cruz sc-56) and PLA probes (anti-mouse MINUS - DUO92004 and anti-rabbit PLUS - DUO92002). Then, cells were incubated with Ligase for 30 minutes (37°C) and Polymerase for signal amplification for 100 minutes (37°C) (Duolink^®^ In Situ Orange - DUO92102). For negative controls, cells were stained, missing one of the primary antibodies, or both. After all PLA protocol steps, cells were stained with anti-FLAG 1:400 and anti-rabbit Alexa Fluor 488 1:1,000 (Invitrogen - A-11008) to identify transfected cells. Nuclei were stained with DAPI. After the respective washes, slides were mounted with Fluoromount-G (Invitrogen). The contrast was evenly enhanced in all presented images for better visualization of PLA dots.

### Confocal microscopy

For co-localization analysis, images were taken with a Zeiss LSM 780 NLO confocal microscope coupled to a HeNe (543 nm), an Argon (488 nm) and a Diode (405 nm) lasers (Core Facility for Scientific Research – University of Sao Paulo (CEFAP-USP)). Images were acquired with an objective α Plan-Apochromat 100x/NA 1.46 in oil immersion. Fluorescent signal was detected on a 32 channel GaAsP QUASAR detector with the following parameters: Alexa Fluor 594 (578 – 692 nm), Alexa Fluor 488 (491 – 587 nm), DAPI (412 – 491 nm). Pinhole was set to 1 airy unit, and z-stacks were taken with intervals of 340 nm. Presented images show a single representative z-stack. Overlap in signal between different channels was measured with Plot Profile plugin on ImageJ. Images were contrast-enhanced for better visualization.

### Western blotting

The proteins were collected from Vero E6 and HEK293 cells using a cell lysis buffer (50 mM Tris-Cl pH 7.5; 150 mM NaCl, 1 mM EDTA, 1% Triton x-100, and protease and phosphatase inhibitor cocktail). To obtain the lysates, cells were maintained with lysis buffer for 15 minutes on ice and centrifuged at 12,000 × g for 10 minutes. Samples containing total protein were separated by SDS-PAGE and transferred to nitrocellulose membranes. Membranes were blocked for one hour at room temperature (RT) with 5% non-fat powdered milk dissolved in TBS-Tween-20 (TBS-T) (50 mM Tris-Cl, pH 7.5; 150 mM NaCl; 0.1% Tween-20), incubated overnight (4°C) with primary antibodies and with secondary antibodies against mouse or rabbit IgG conjugated with peroxidase (Amersham) for one hour at RT. The membranes were washed three times with TBS-T, incubated with the SignalFire ECL Reagent (Cell Signaling Technology) for protein bands visualization. The band’s densitometry was performed using ImageJ software v1.53 (National Institutes of Health). The following antibodies were used: anti-PCNA (#2586); anti-γH2AX (#9718); anti-β-actin (#4967) (Cell Signaling), anti-α-tubulin (CP06) (Calbiochem)

### Immunohistochemistry

Lung tissue sections, obtained via Minimally Invasive Autopsies, (COVID-19 case and a control lung tissue of a non-smoker subject) were stained in silanized slides (Sigma Chemical Co. St. Louis, Missouri, EUA). Briefly, the histological sections were deparaffinized in xylene, rehydrated through a graded series of ethanol, and kept in a tri-phosphate buffer (TBS) pH 7.4. After deparaffinization, slides were hydrated for five minutes in graded alcohol series (100%, 95%, 70%, and H_2_O) and incubated in Tris-EDTA for 50 minutes at 95°C pH 9.0. The blockage of the endogenous peroxidases was done by hydrogen peroxide 10v (3 %)

The slides were then incubated with Ab mix (TBS (1%), BSA (4%), and Tween20 (0.02%)) for 20 minutes at RT followed by treatment overnight at 4 °C with anti-γ-H2AX (Cell Signaling #9718S) diluted 1:500 in Ab mix or anti-PCNA diluted 1:1000 (DAKO #M0879). The slides were then washed twice in TBS (1%) and finally incubated for one hour at RT with 1:500 goat anti-rabbit/mouse HRP polymer detection kit – ImmunoHistoprobe Plus (ADVANCED-BIOSYSTEMS) for 30 minutes at 37°C). The diaminobenzidine (DAB) was used as a chromogen (Sigma-Aldrich Chemie, Steinheim, Germany). Finally, the counterstaining was done with Harris Hematoxylin (Merck, Darmstadt, Germany), and the slides were mounted with a coverslip in Permount (Fischer #SP15-500). The use of this material for research purposes has been previously approved by the Institutional Ethical Board CAAE #30364720.0.0000.0068).

### Anti-viral *in vitro* efficacy assay

A plaque assay was performed as described in Kashyap *et al*. 2021 ^42^. Briefly, after seeding, Vero E6 (8 × 10^5^ cells per well, 6-well plates) were incubated overnight and infected with 200 PFU per well. PCNA-I1 (Cayman #20454) or Verdinexor (Cayman #26171) were added at final concentrations of 0.5 μM and 0.1 μM, ^41^ in overlays composed of DMEM supplemented with 10% FBS plus carboxymethylcellulose sodium salt (Sigma-Aldrich #C5678) 2%, immediately post-viral adsorption. After four days, cells were fixed with formaldehyde and stained with crystal violet to count plaques.

### Statistical analysis

Data are presented as mean ± Standard Deviation (SD). Statistical analysis of the data was performed by Student’s t-test or ANOVA. P-values of ≤0.05 were considered statistically significant. All experiments were performed in duplicate. The odds ratio was calculated to verify if the translocation of the PCNA to the cytoplasm was dependent on the M transfection. A total of 147 cells were analyzed. A value of odds ratio greater than one indicates that the event observed is dependent on the object analyzed.

## Acknowledgements

We acknowledge the students that helped EPZ during the cell lines infection in the Biosafety Level 3 Laboratory (NB3) and all others that were our backup outside of the lab.

## Author contributions statement

Conceptualization, FMS and AMV; methodology, EPZ, ICBP, MCSM, APM, MBS, OBS, MMG, KBS, MRA; software, MBS; validation, EPZ, ICBP, MCSM, APM, MBS, OBS and MMG; formal analysis, EPZ, FMS, HMS and AMV; investigation, EPZ and FMS; resources, FMS, JLPM, HMS, TM, MD, PHNS, and AMV; data curation, EPZ, MMCS, MBS, FMS, HMS, and AMV; writing original draft preparation, EPZ; writing—review and editing, EPZ, FMS, JLPM, and AMV; visualization, EPZ, ICBP, MCSM, APM, MBS, OBS, MMG, KBS, MRA, DATT, PLAP; supervision, FMS, JLPM, HMS, and AMV; project administration, FMS and JLPM; funding acquisition, FMS and JLPM. All authors have read and agreed to the published version of the manuscript.

