## Supplemental Figures for "Characterization of the interaction between SARS-CoV-2 Membrane Protein and Proliferating Cell Nuclear Antigen (PCNA) as a Potential Therapeutic Target"

**CONTENTS**

**Supplementary Figures: 1-4**

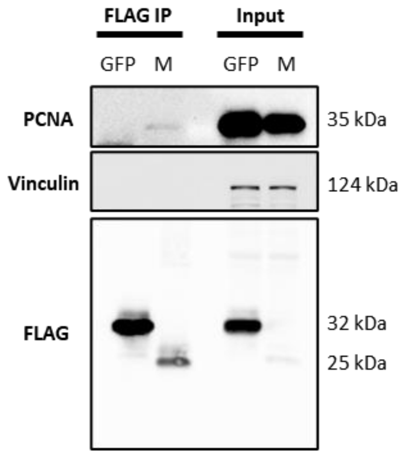


**Supplementary Figure S1.** Confirmation of M interaction with PCNA in another immunoprecipitation assay.

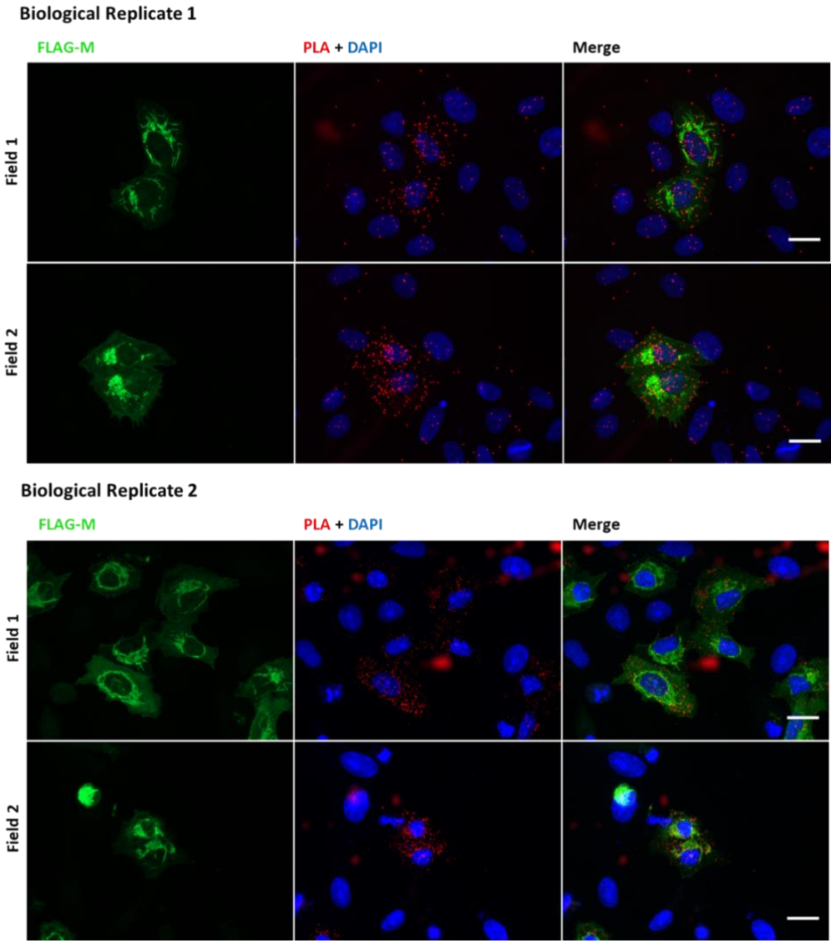


**Supplementary Figure S2. Representative fields of PLA positive signal.** The figure shows two different fields of non-cropped images for two independent biological replicates. All images were taken at 63× magnification with a ZEISS Axio Vert.A1 microscope. Scale bars 20 µm.

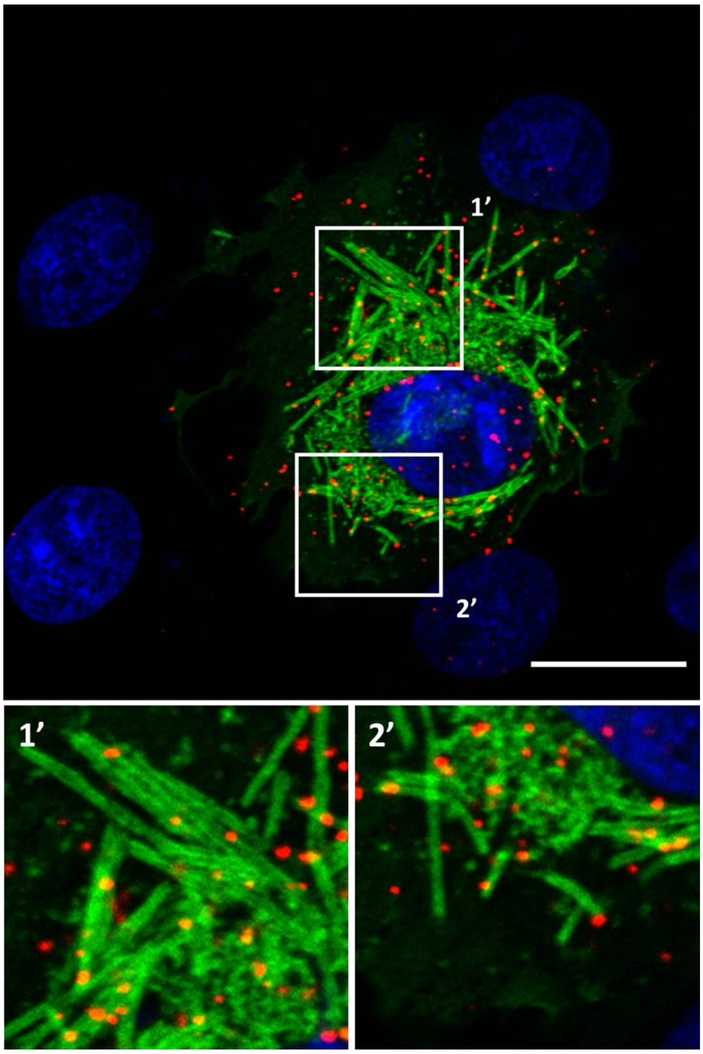


**Supplementary Figure S3. Detection of FLAG-M/PCNA interaction by proximity ligation assay.** Positive PLA cells were analyzed by confocal immunofluorescence to investigate proximity between PLA signal and FLAG-M membranous structures. Panels 1’ and 2’ represent zoomed areas indicated by framed regions in merge panel, showing PLA dots colocalizing with FLAG-M. The figure shows a single plane from a z-stacked image. Images were taken at 100× magnification with a Zeiss LSM-780-NLO microscope. Scale bars 20 µm.

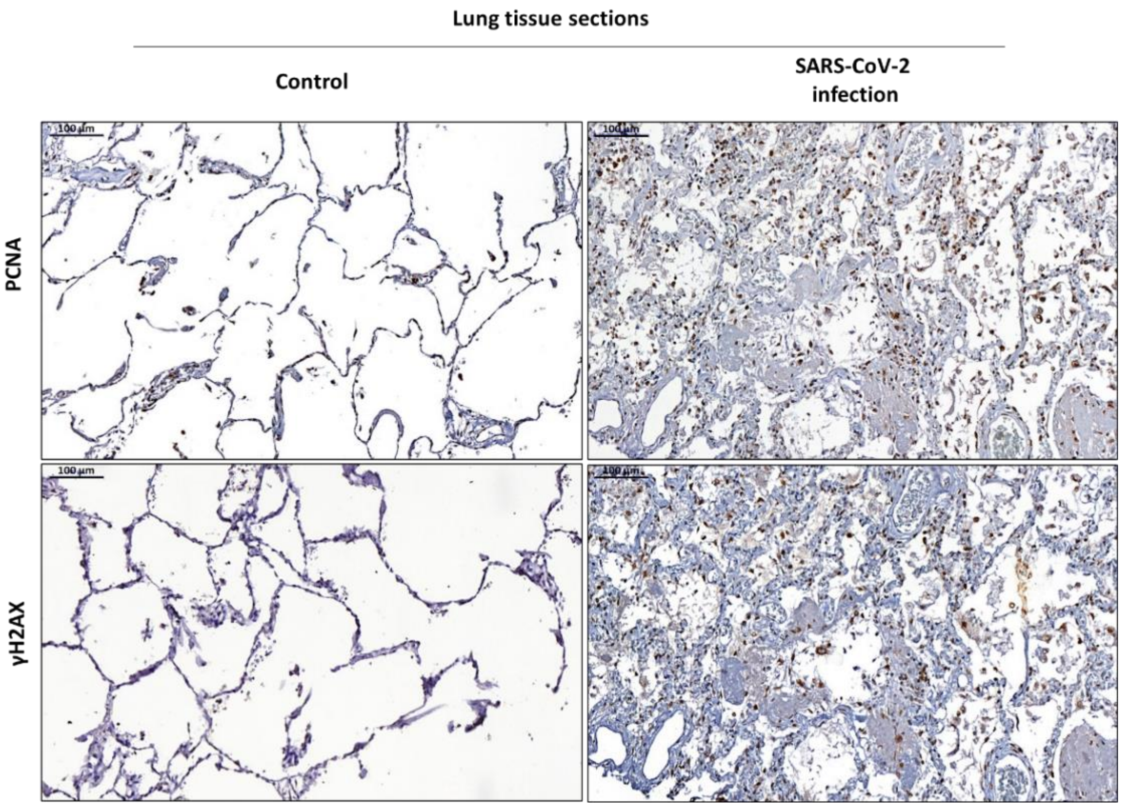


**Supplementary Figure S4. Immunohistochemistry of lung sections from one control case and one COVID-19 patient immunostained with PCNA and γH2AX.** Immunohistochemical positivity (brown color) for PCNA and γH2AX proteins are present in cells along the alveolar epithelium, with a higher density of positively stained cells for both markers in the COVID-19 case (right panels). This data is representative of one independent experiment.
